# A simple workflow to identify novel Small Linear Motif (SLiM)-mediated interactions with AlphaFold

**DOI:** 10.1101/2025.06.04.657817

**Authors:** M. Veinstein, V. Janssens, B.I. Iorga, R. Helaers, T. Michiels, F. Sorgeloos

## Abstract

Short linear motifs (SLiMs) are highly compact interaction modules embedded within disordered protein regions and are increasingly recognized for their central role in maintaining cellular homeostasis. Due to their small size, degeneracy and transient binding, SLiMs remain difficult to detect both experimentally and computationally. Here, we show that AlphaFold, used via ColabFold, offers a practical and accessible alternative for *in-silico* SLiM discovery.

Unlike previous studies focused on structural accuracy, we evaluated AlphaFold’s capacity to reveal SLiMs independently of model quality. To this end, we benchmarked several scoring metrics and showed that AlphaFold2 combined with MiniPAE yields the best performance, outperforming AlphaFold3 in this context.

Building on these findings, we also provide a streamlined and cost-effective workflow for SLiM prediction requiring no installation or local computation. To overcome challenges associated with SLiM validation, we also introduce a highly sensitive detection method based on proximity labeling in living cells. This workflow was used to predict the occurrence of SLiMs that mediate binding to ribosomal protein S6 kinase A3 (RPS6KA3 or RSK2).

By leveraging Colabfold and MiniPAE available through Colab notebooks, our approach provides a scalable and widely accessible strategy for identifying functional SLiMs in proteins of interest.

MiniPAE can be accessed at https://github.com/martinovein/MiniPAE

**Short description:** Martin Veinstein is a PhD student in Biomedical Sciences at the de Duve Institute, UCLouvain, Belgium. He specializes in Small Linear Motifs (SLiMs) in the context of host–virus interactions and has developed strong expertise in bioinformatics, structural biology, and predictive modeling.

Victor J is a unfergradiate student at the ECAM Brussels Engineering School, Haute Ecole “ICHEC-ECAM-ISFSC”, Brussels, Belgium. His activities span form September to November 2023.

B.I. Iorga is a CNRS Research Director at the Institut de Chimie des Substances Naturelles in Gif-sur-Yvette, France. His research focuses among others on methodological developments in molecular modeling and the *in-silico* prediction of antibiotic resistance using machine learning and deep learning approaches.

Raphael Helaers is a Senior Investigator and leads bioinformatics infrastructure at the de Duve Institute, UCLouvain, Belgium. He has developed strong expertise in next-generation sequencing and software development, along with a deep interest in biology, genetics, and evolution.

Thomas Michiels is a Full Professor and researcher at the de Duve Institute, UCLouvain, Belgium. His research focuses on virus-mediated subversion of the innate immune response.

Frederic Sorgeloos is an adjunct Professor at the INRS, Laval, Canada. He currently focuses on the subversion of cellular homeostasis through small linear peptides encoded by viral and bacterial pathogens.

**Short abstract:** Various AlphaFold2/3 scoring metrics were systematically benchmarked for their ability to detect Small Linear Motifs (SLiMs)

Based on this evaluation, a user-friendly and cost-effective *in-silico* workflow is proposed to identify novel SLiMs-targeting proteins

The utility of this workflow is demonstrated through the prediction of previously uncharacterized SLiMs interacting with RSK kinases.

A sensitive *in-vitro* assay is proposed to streamline the validation of low-affinity SLiM-target interactions.

Together, our workflow and associated validation assay offer an integrated pipeline for the discovery and validation of SLiM-mediated protein-protein interactions.

## Introduction

Short linear motifs (SLiMs) are short regions of 3-10 amino acids embedded within intrinsically disordered regions (IDRs) that mediate protein–protein interactions (PPIs). These interactions generally occur between a SLiM and a structured protein domain and exhibit low-to-mid micromolar affinities, making them well suited for dynamic and reversible interactions that regulate cell signaling (1). SLiMs control a wide range of biological processes and are catalogued in resources such as the Eukaryotic Linear Motif (ELM) database, which currently nemocontains around 1,500 annotated human SLiMs (2). However, given the estimated tens of thousands of human SLiMs, the vast majority of them remain undiscovered (1).

Despite their importance, SLiMs are notoriously challenging to detect directly due to their small size and weak binding affinities. Alternative *in-vitro* methods have been developed to improve detection sensitivity (3). However, these approaches remain labor-intensive, difficult to scale and still face intrinsic affinity limitations. As such, computational approaches have emerged as powerful complementary tools to accelerate SLiM discovery.

Despite not being specifically designed for this purpose, AlphaFold2 (AF2) has shown an unexpected ability to model SLiM-mediated interactions. An early study demonstrated that AF2 could accurately predict the structure of interactions between IDRs and their structured binding domains, even for protein pairs entirely absent from its training set (4).

Subsequent benchmarking by Lee *et al.* provided a systematic comparison of different AlphaFold-based metrics for SLiM–domain interactions (5). Notably, they were among the first to apply AlphaFold at scale to screen for SLiM-mediated PPIs across an entire signaling network. Their results, however, revealed limited specificity, highlighting the need for optimized scoring strategies when applying AlphaFold in unbiased screenings.

More recently, Omidi *et al.* conducted a comprehensive analysis of AlphaFold2’s ability to model various IDR-mediated interactions including SLiMs, Molecular Recognition Features (MoRFs), and others (6). Importantly, they introduced two new residue-level metrics: MinD, which evaluates the minimum distance between residues across interfaces, and MiniPAE, derived from AlphaFold’s Predicted Aligned Error (PAE) maps, to identify potential interaction regions within full-length IDRs.

In this study, we moved beyond evaluating AlphaFold’s ability to produce accurate structures and instead assessed its capacity to identify SLiMs. To this end, we systematically compared the main AlphaFold-derived scoring functions used for SLiM discovery and extended this evaluation to include AlphaFold3 (7). This analysis led to a streamlined and accessible workflow for SLiM discovery, which we have made available through a user-friendly Colab notebook. We illustrated the power of this approach with a case study focused on identifying RSK-binding SLiMs. To complement our *in-silico* predictions and address the challenge of low-affinity interactions, we further introduced a sensitive proximity labeling method, enabling functional validation, in cells, of candidate SLiMs predicted by our pipeline.

## Results

### 1. Benchmark SLiM datasets

While recent publications have primarily evaluated AlphaFold’s ability to generate accurate structural models of SLiM-mediated interactions, our focus here is distinct: we assessed AlphaFold’s capacity to identify SLiMs within binary interaction predictions, irrespective of the structural quality of the predicted complex.

To this end, we constructed a benchmark dataset derived from the Eukaryotic Linear Motifs (ELM) database, a manually curated repository of experimentally validated SLiMs. Given the high redundancy of SLiMs, the ELM database groups analogous motifs under a common classification, termed ELM_Class. Among the 161 ELM_Classes annotated in humans, we selected 121 representative instances, each corresponding to a distinct ELM_Class, to serve as our positive dataset, hereafter referred to as the ELM_dataset (see Methods, Figure 1A, Table S1).

**Figure 1:**
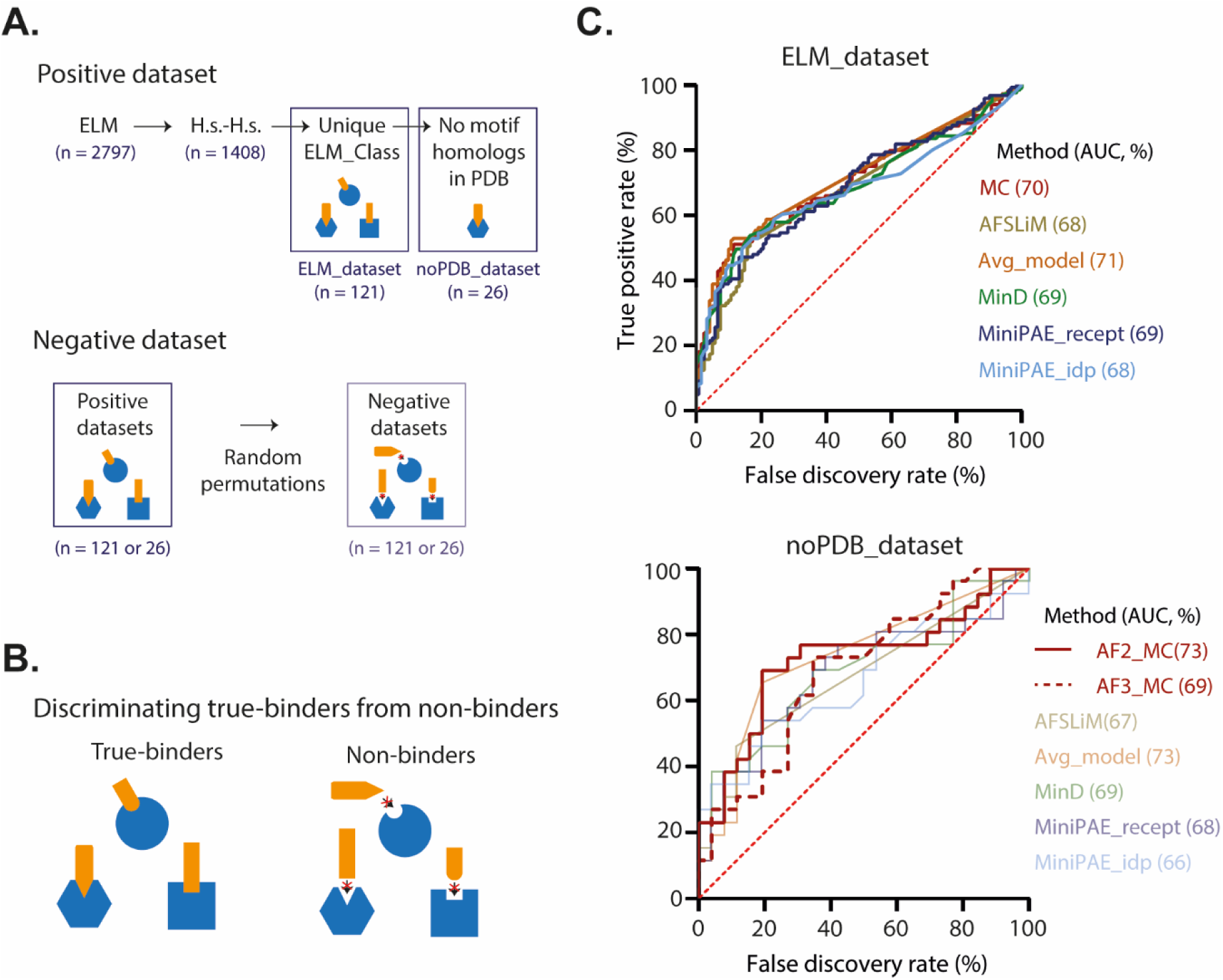
Benchmarking of AlphaFold-based metrics for SLiM-mediated interaction detection. **A.** Construction of benchmark datasets. A non-redundant set of 121 human SLiM–partner pairs, each representing a unique ELM_Class, was extracted from the ELM database resulting in the ELM_dataset (Table S1). A subset of 26 instances lacking structural homologs in the Protein Data Bank (PDB) constituted the noPDB_dataset. Negative datasets were generated by randomly permuting the interaction partners within each positive dataset. **B.** Overview of the first subproblem in SLiM prediction using AlphaFold. **C.** Receiver Operating Characteristic (ROC) curves comparing true-binders to randomly permuted non-binders, using indicated datasets. Results are shown for AlphaFold2 (AF2, solid lines) and AlphaFold3 (AF3, dashed lines). Five scoring metrics are compared: Model Confidence (MC), AlphaSLiM (AFSLiM), average_model from Schmid *et al*. (9) (Avg_model), and the minD and miniPAE metrics from Omidi *et al*. Area Under the Curve (AUC) values are indicated between parentheses for each metric.

Unlike structure-centric benchmarks, our evaluation does not require the existence of homologous complexes in the Protein Data Bank (PDB). This allowed us to form a subdataset of 26 SLiM–partner pairs which have no available structural homologs. This dataset was named noPDB_dataset (Figure 1A, Table S1).

To accurately assess AlphaFold’s predictive performance, we also established a negative control set composed of likely negative pairs. This was achieved by random permutations of binding partners within both positive datasets, following the approach of Lee *et al.* (5) (Figure 1B, Table S1).

### 2. Comparison of different AlphaFold2/3 metrics to discriminate SLiM-binders from non-binders

AlphaFold’s utility in SLiM discovery can be conceptually divided into two challenges. The first is binary **classification**—determining whether a SLiM mediates binding to a partner at all. The second is **localization**—identifying the specific region that functions as a SLiM. For those tasks different AlphaFold-based metrics can be used.

Among those, pLDDT has garnered considerable attention as a tool for localizing SLiMs (8). Notably, Lee *et al.* (5) proposed pLDDT as the most robust metric for SLiM detection, and Omidi *et al.* (6) identified it as the best discriminator between binding IDRs and decoy regions. However, despite this enthusiasm, pLDDT suffers from a fundamental limitation: it tends to increase not only in disordered regions upon binding—the signal of interest—but also in pre-structured regions, regardless of their direct involvement in protein-protein interactions. This dual behavior blurs interpretation and undermines its specificity for SLiM identification, a limitation also reflected in the findings of Omidi *et al*.

To address this limitation, we developed AlphaSLiM, a new scoring metrics that quantifies the change in per-residue pLDDT between the unbound and bound states of an IDR (see Methods). We compared AlphaSLiM to four other AlphaFold-based metrics: the Model Confidence (MC) output by AlphaFold, the average_model proposed by Schmid *et al.* (9), and the minD and miniPAE scores proposed by Omidi *et al*.

Using our ELM_dataset, we first evaluated these metrics for their ability to distinguish true SLiM-binders from non-binders. All metrics—AlphaSLiM, MC, average_model, minD, and miniPAE—achieved comparable performance, with ROC AUC values around 70% (±3%) (Figure 1C, Table S1) suggesting that AlphaFold-based scoring functions can moderately discriminate binders from non-binders. The performance is even more limited in practical settings: at a false discovery rate (FDR) of 0, where the true positive rate (TPR) dropped to near 10%. Importantly, when we repeated the evaluation using the noPDB_dataset, results remained consistent (AUC = 70% ± 4%, Figure 1C, Table S1), indicating that the observed performance of AlphaFold-based metrics in distinguishing binders from non-binders did not arise from structural biases linked to PDB templates.

To evaluate whether newer model architectures improve SLiM detection, we compared AlphaFold2 and AlphaFold3. To avoid biases stemming from differences in training data, especially with respect to known structural templates, we restricted this comparison to the **noPDB_dataset**. The results revealed comparable performance across AlphaFold2 and AlphaFold3 metrics in terms of binder versus non-binder discrimination, suggesting that despite its architectural advances, AlphaFold3 model confidence (MC) does not provide a clear advantage for SLiM binders from non-binders discrimination (Figure 1C, Table S1).

### 3. Comparison of different AlphaFold2/3 metrics to identify new SLiMs within full-length proteins

We next assessed AlphaFold’s ability to localize a binding SLiM within full-length proteins (Figure 2A). For this, we compared the per-residue score within the SLiM region when paired with its true partner to the score within the same region when paired with a random non-binder (Figures 2B–C). With our ELM_dataset, all three metrics showed similar overall performances (AUC ∼78%, Figure 2B), but miniPAE outperformed the others at FDR = 0. This advantage disappeared with the noPDB_dataset (Figure 2C–D; Supplementary Figure S1A), suggesting that miniPAE’s precision is driven by structural similarity to known templates.

**Figure 2:**
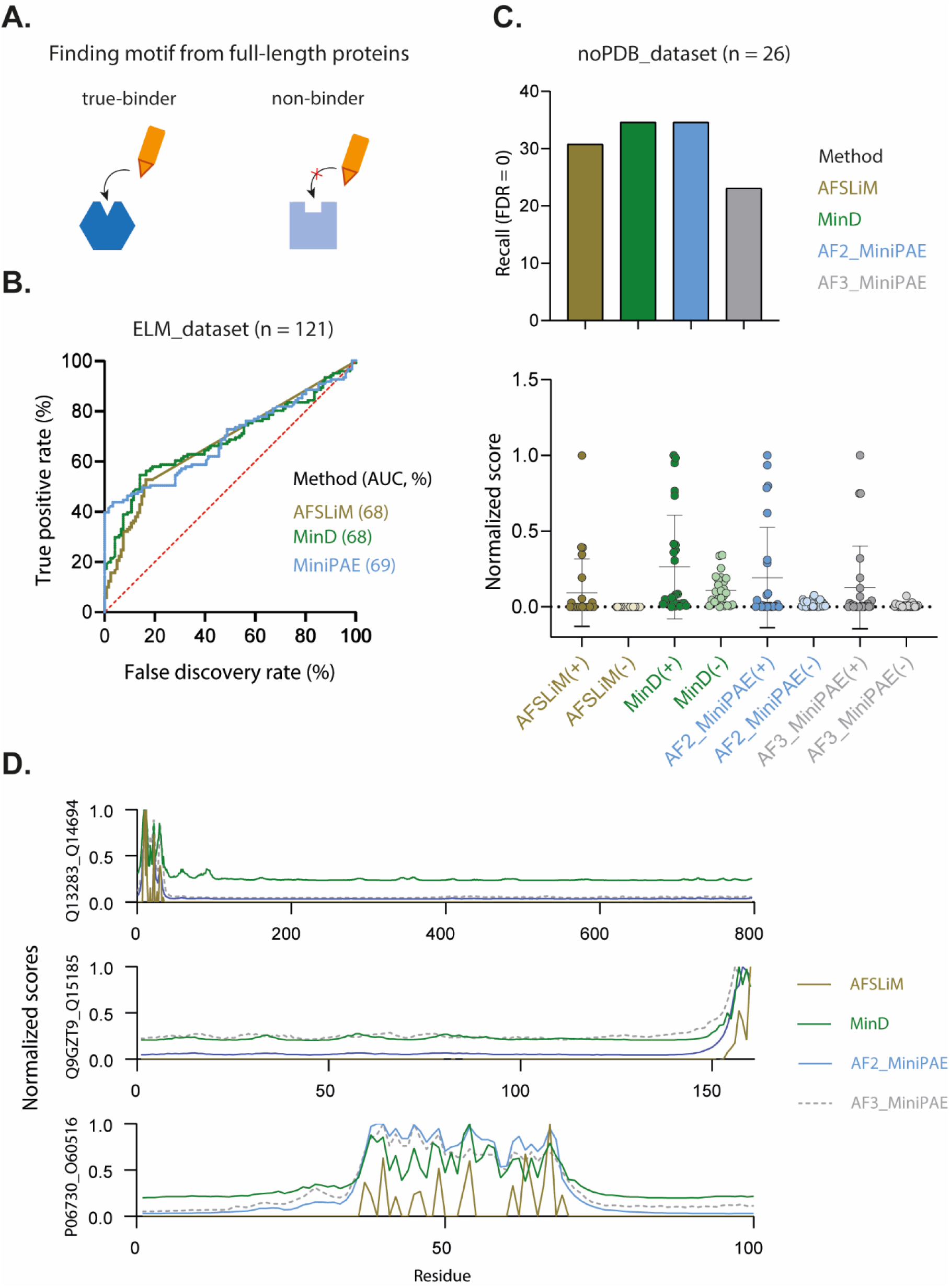
Using AlphaFold metrics to localize SLiMs within full-length proteins. **A.** Schematic of the analysis: scores within the motif region when paired with its true-binder are compared to the score within the same region when paired with a random non-binder. **B.** ROC curves for the ELM_dataset. **C. Top**: True Positive Rate (TPR, %) at FDR = 0. **Bottom**: Distribution of scores in the motif region for true-binders (+) vs non-binders (-) on a 0,1 scale (see method). **D.** Examples of per-residue score profiles for indicated protein pairs.

The large overlap between true/false motif calls from AlphaSLiM, minD, and miniPAE (Supplementary Figure S1B) suggests that combining these metrics offers little added benefit. Additionally, although shorter prediction lengths are known to improve structural accuracy (4,5), we found no correlation between prediction length and motif detection (Spearman ρ = 0.03 and –0.17 for ELM_dataset and noPDB_dataset, respectively; Supplementary Figure S1C).

Finally, since both AlphaFold2 and AlphaFold3 output PAE scores, we used the noPDB_dataset to compare their ability to localize SLiMs within proteins. AlphaFold3 showed slightly lower performance than AlphaFold2 in this task (Figure 2C, Supplementary Figure S1D; Table S1).

### 4. AlphaFold-based workflow to identify new SLiMs binding a protein of interest

Building on our benchmarking results, we propose a step-by-step workflow to discover new SLiMs binding a specific protein (Figure 3), while highlighting common pitfalls in such AlphaFold screenings. The first critical decision lies in selecting candidate protein partners for binary prediction. A highly ambitious approach would be to perform blind screening against entire proteomes. However, AlphaFold-based metrics are not well-suited for this approach as they perform poorly in blind screenings where non-binders vastly outnumber true-binders, as shown by Schmid *et al.* (9) and confirmed with our results in Figure 1C. To mitigate this, we recommend selecting partners from curated interaction databases such as BioGRID or IntAct, or from in-house datasets. Given that AlphaFold2 slightly outperforms AlphaFold3, and is readily accessible, we suggest its use for most cases.

**Figure 3:**
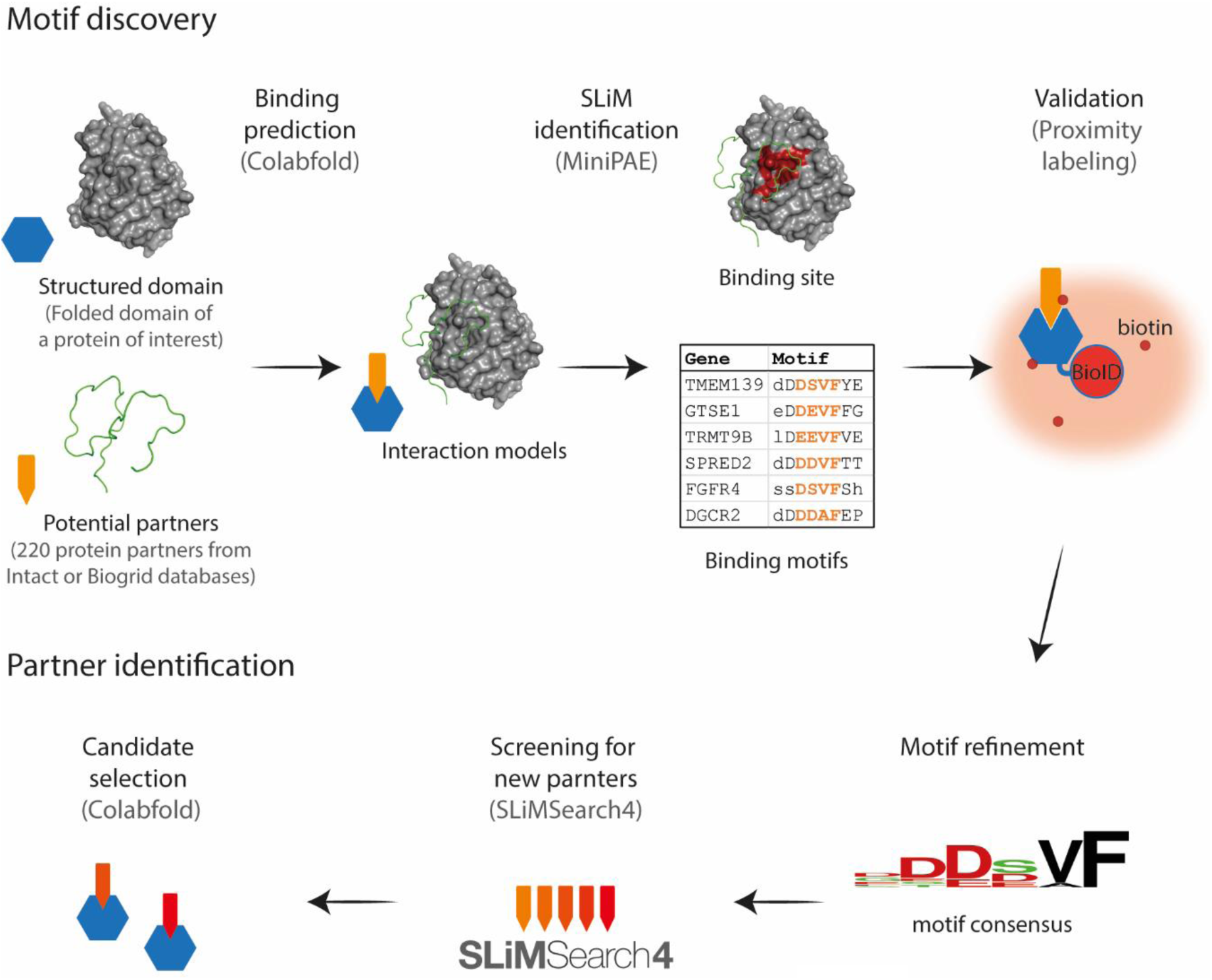
An AlphaFold-based workflow to identify new SLiMs. ColabFold is used to perform binary binding predictions between a protein (or domain) of interest and selected candidates from interaction databases or in-house datasets. The MiniPAE script extracts high-confidence SLiM predictions, which can be experimentally validated using proximity labeling strategy owing to their typically low affinity. Finally, this motif knowledge can be used to discover new interactants of your protein of interest using SLiMSearch4. All *in-silico* tools can be accessed via colab notebooks or webservers: Colabfold: https://colab.research.google.com/github/sokrypton/ColabFold/blob/main/AlphaFold2.ipynb#scrollTo=kOblAo-xetgx AF2_MiniPAE: https://colab.research.google.com/drive/1ZAj5gQOP-usd8yAK5uhoLsLH4WPAKxMG SLiMsearch4: https://slim.icr.ac.uk/tools/slimsearch/input

To streamline SLiM retrieval, we adapted the MiniPAE script from Omidi *et al.* (6) to analyze AlphaFold2 output files, returning MiniPAE scores, motif sequences, and their coordinates. Since experimental validation is often more limiting than prediction time, we suggest using a MiniPAE threshold of 4, which corresponds to an FDR of 0 on the ELM_dataset (Figure 2B).

With regard to SLiMs, validating them remains a significant challenge given the low affinity interaction that they often mediate (see Alexa *et al.,* review). To mitigate this, we propose a proximity labeling assay as a low-cost, sensitive method to detect interactions in living cells. This involves expressing the predicted SLiM expressed as a fusion protein (see Methods) and assessing its proximity enrichment relative to the target protein. Once validated, interacting motifs can be aligned to infer the minimal binding consensus. While identifying the interaction motif of a known partner is informative, the ultimate goal is often to discover new binders. This can be achieved using tools such as SLiMSearch 4 (10), which scans the proteomes for additional motif matches. Depending on the specificity of the consensus, this may yield a narrow or broad spectrum of candidates that can be further refined using AlphaFold to evaluate whether the motifs adopt a compatible binding interface *in-silico*.

### 5. Illustration of the use of our workflow to find hRSK2-binding SLiMs

To illustrate the practical application of our SLiM discovery workflow, we used it to identify novel motifs binding human RSK2 (hRSK2), a Ser/Thr kinase broadly expressed across tissues and involved in numerous biological processes (11,12). Mutations in RSK2 are known to cause Coffin-Lowry syndrome (13) and have been implicated in several other disorders (11), underscoring the importance of understanding how RSK2 selectively engages its partners through SLiMs.

As outlined in the previous section (Figure 3), we started by selecting candidate RSK2-binding proteins. For this, we queried the BioGRID and IntAct interaction databases, and, recognizing RSK2’s role as a kinase, we also included all known substrates listed for RSK2 in PhosphoSitePlus. This resulted in a list of 220 proteins with less than 1000 amino acids (longer proteins were not considered), which were then individually modeled in binary complexes with RSK2 using AlphaFold2. The resulting structures were analyzed using our adapted MiniPAE script, and motifs were filtered using a MiniPAE threshold of 4, as established in the previous section. This analysis yielded 21 predicted SLiMs from 20 proteins.

Predicted SLiMs clustered into two distinct binding classes. The first group consisted of 10 “DDVF-like” motifs that engaged a region centered on the KAKLGM loop of RSK2—a surface previously implicated in regulatory interactions and viral hijacking (14,15). The second group, comprising 11 “substrate motifs”, bound the catalytic cleft of RSK2 and resembled canonical substrate motifs (Figure 4A, Table S2).

**Figure 4.**
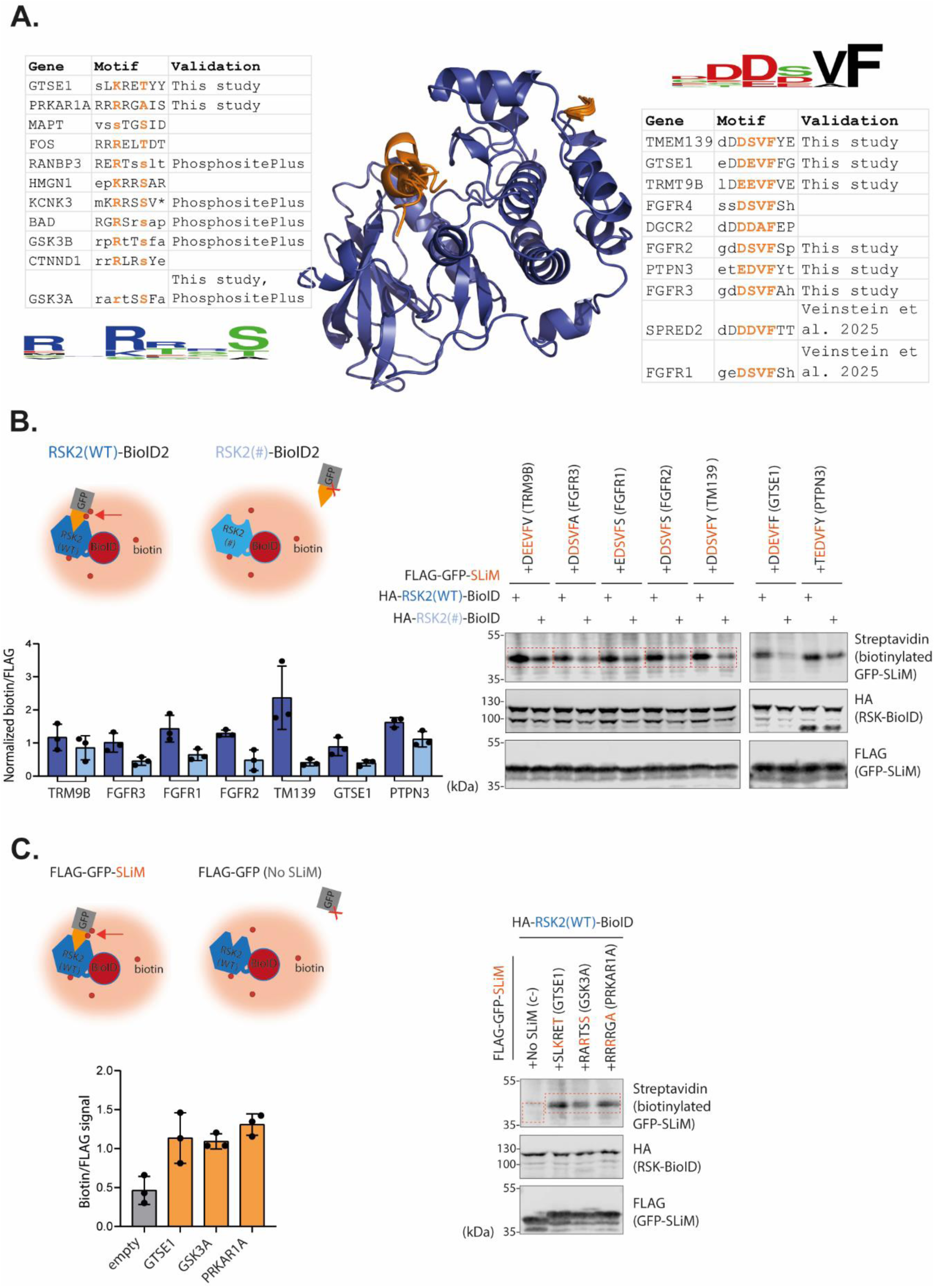
Discovery and validation of hRSK2-binding SLiMs using our AlphaFold-based workflow. **A.** AlphaFold2 predictions show two classes of SLiMs (orange) binding to hRSK2 (blue): substrate motifs (left) and DDVF-like motifs (right), along with their associated consensus logos. **B–C.** Experimental validation of SLiM–RSK2 proximity using a BioID2 proximity labeling. **B:** DDVF-like motifs were tested with wild-type or KSEPPY-mutant RSK2. **C:** Substrate motifs were tested with wild-type RSK2 and compared to the empty construct. Schematics (top), representative immunoblots (right), and quantifications (left) confirm motif-specific proximity labelling (n = 3).

To experimentally validate these predictions, we selected a subset of representative SLiMs from each class. Among the DDVF-like motifs, two had previously been shown to bind RSK1, which makes sense given that the closely related RSK1 and RSK2 share the same KAKLGM loop (14,15). We focused on six new DDVF-like motifs for validation. Given the low affinity of SLiM interactions, we used a proximity labeling strategy involving BioID2 fused to RSK2. Each predicted SLiM was cloned at the C-terminus of a FLAG-GFP construct, and biotinylation of this protein was assessed in cells co-expressing the fusion between BioID2 and either wild-type RSK2 or a KAKLGM-to-KSEPPY mutant previously shown to reduce DDVF-mediated interactions. For all six SLiMs examined, the KAKLGM mutation in RSK2 reduced biotinylation of the FLAG-GFP-SLiM constructs, confirming RSK2 KAKLGM loop-dependent interactions. However, the extent of biotinylation enrichment (WT-RSK2/mutant RSK2) varied, with TMEM139 showing the most pronounced motif-specific proximity labeling, while TRM9B and PTPN3 exhibited smaller intensity differences.

For the “substrate SLiMs”, five of the eleven predicted motifs were already annotated as RSK2 phosphorylation sites according to PhosphositePlus. We selected two uncharacterized motifs from the remaining six. Validating these was more challenging: substrate motifs are not expected to form stable interactions with kinases due to the transient nature of catalysis, and there are no well-characterized mutations that abolish substrate “binding” without affecting kinase function. Therefore, we assessed proximity by comparing the biotinylation levels of FLAG-GFP-SLiM constructs to those of FLAG-GFP alone. As shown in Figure 5, the addition of the motif led to increased biotinylation, suggesting that our predicted substrate motifs were indeed contributing to the “interaction” with RSK2. Notably, these motifs matched the known consensus sequence for RSK2 substrates, as curated in PhosphoSitePlus, reinforcing the validity of our predictions.

**Figure 5.**
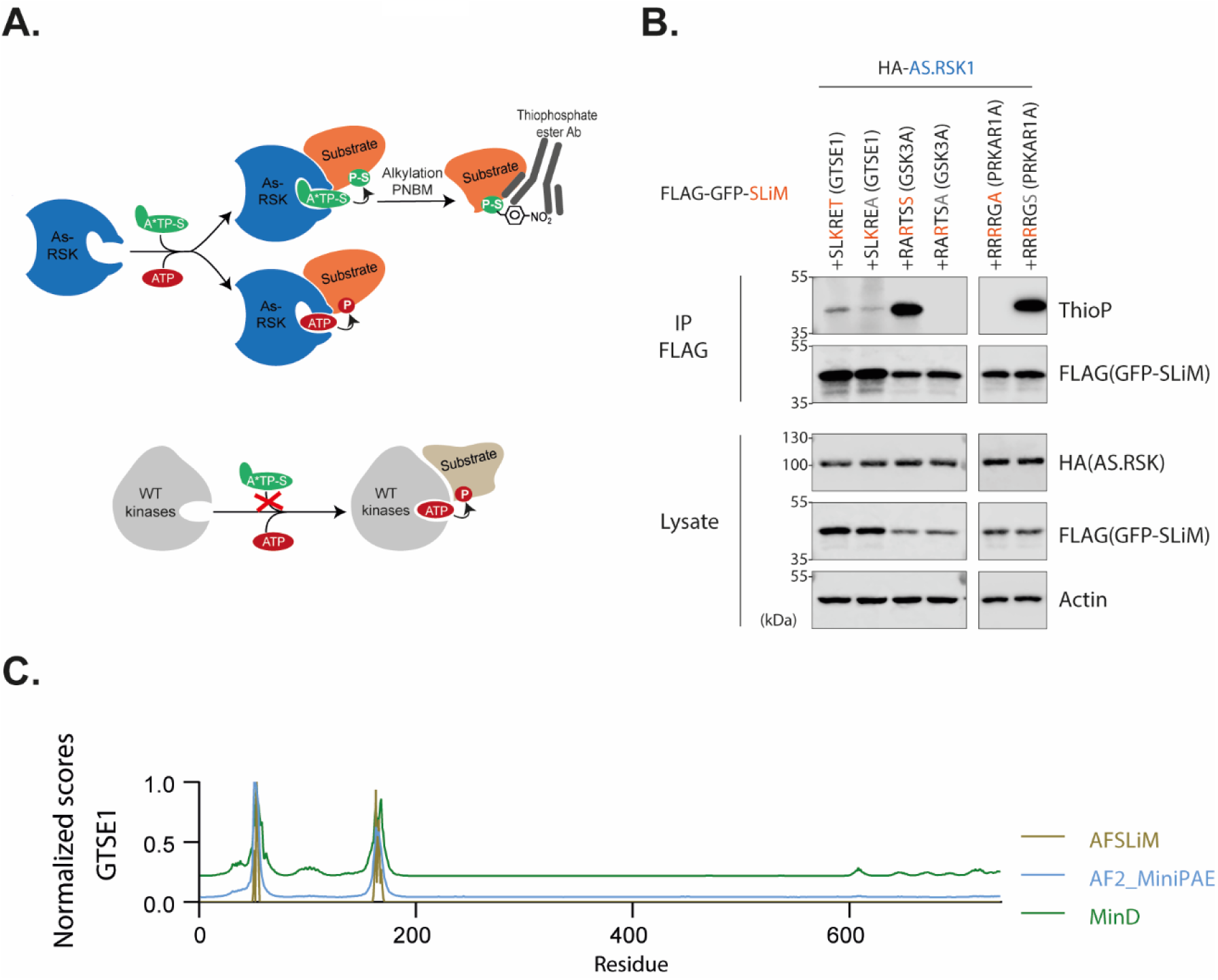
RSK1-dependent phosphorylation of predicted substrate SLiMs using an analog-sensitive kinase system. **A.** Schematic representative of the analog-sensitive kinase system (adapted from Lizcano-Perret *et al*. (16). The AS kinase incorporates a thiophosphate from N6-Bn-ATP-γ-S (A*TP-S) into substrates, enabling subsequent alkylation with PNBM and detection via a thiophosphate ester-specific antibody. **B.** Representative immunoblots showing thiophosphorylation of immunoprecipitated FLAG-GFP-SLiM proteins harboring predicted substrate motifs. Constructs were transfected in BLP32 cells expressing HA-tagged AS-RSK1. Wild-type motifs (orange) exhibit greater RSK1-dependent thiophosphorylation compared to their mutant counterparts (grey) (n = 2).

While these results showed that substrate motifs conferred proximity to RSK2, this alone could not confirm whether the kinase actually phosphorylated them. To address this, we used an analog-sensitive (AS) kinase system previously developed for RSK1 by Lizcano-Perret *et al*. (16), taking advantage of the high sequence conservation across RSK paralogs (12). This system enabled the specific detection of phosphorylation events mediated by an analog sensitive RSK (Figure 5B). For both GTSE1 and GSK3A substrate motifs, phosphorylation of the wild-type SLiM was significantly higher than that of the non-phosphorylatable serine-to-alanine (S>A) mutant, confirming RSK2-dependent phosphorylation (Figure 5B). Notably, the phosphorylation signal for GTSE1 was markedly weaker than for GSK3A, consistent with the fact GSK3A has a canonical substrate motif while GTSE1 does not (12). We hypothesize that GTSE1’s dual DDVF and substrate motifs may compensate for its non-canonical sequence, facilitating phosphorylation in the context of the full-length protein. In the case of a predicted pseudosubstrate motif in PRKAR1A that naturally contains a non-phosphorylatable alanine at the phospho-acceptor site, we did the converse and introduced an alanine-to-serine mutation and observed phosphorylation of the mutant, further supporting the accuracy of our prediction.

Altogether, this experimental setup allowed to confirm, to various degrees, all eight tested SLiMs, supporting the use of a MiniPAE threshold of 4 for predictive screening. Regarding substrate motifs, our results suggest that AlphaFold predictions can extend beyond structural binding to inform on enzymatic modification events, consistent with previous suggestions by Lee *et al*. (5) in the context of protease cleavage site prediction.

## Material & Methods

### Benchmark

We downloaded all 2,797 interaction instances from the ELM database and retained only those corresponding to *Homo sapiens–Homo sapiens* interactions. Following the filtering strategy of Bret *et al.* (4), we excluded instances in which the same motif occurred multiple times within a single protein, as this can complicate result interpretations. We also removed a small number of entries for which the reported motif consensus (“regular expression”) could not be located within the corresponding protein sequence. To avoid redundancy, we kept the shortest instance per ELM class. In addition, to reduce prediction time, we restricted both motif-containing proteins and their partners to sequences shorter than 1,000 residues, yielding a final benchmark set of 121 interactions—referred to as the **ELM_dataset** (see Figure 1A)— across 161 unique human ELM classes.

Among these, only 26 interactions had no corresponding structural information available in the PDB according to ELM annotations; this subset is referred to as the **noPDB_dataset**.

For both datasets, we generated a size-matched negative control set by randomly permuting motif-partner pairs while ensuring that no new functional motifs were introduced by chance, as verified using the original motif regular expression (see Figure 1A). All instances are provided in Table S1.

### AlphaFold prediction

Protein complex structures were predicted using AlphaFold2 via ColabFold in multimer V3 mode with default parameters. Predictions using AlphaFold3 were performed using the AlphaFold Server available at https://alphafoldserver.com/.

### MiniPAE and other AlphaFold-based scoring metrics

#### Classical AlphaFold-based metrics

Per-residue pLDDT and Predicted Aligned Error (PAE) matrices were extracted from ColabFold output files. Global confidence metrics (pTM, ipTM) were combined into a Model Confidence score: MC = 0.2×pTM + 0.8×ipTM (5)

#### MiniPAE

For binary complex predictions, AlphaFold outputs a PAE matrix that quantifies the confidence in the relative positioning of different residues. MiniPAE, introduced by Omidi *et al*. (6), represents the lowest (i.e., best) inter-protein PAE value in this matrix. We implemented a simplified MiniPAE script for both AlphaFold2 and AlphaFold3 available here:

https://github.com/martinovein/AF2_MiniPAE

https://github.com/martinovein/AF3_MiniPAE

#### AlphaSLiM score

While pLDDT is commonly employed for SLiM detection, its utility is limited by increased values in both structured regions and disordered regions upon binding. To overcome this limitation, we developed the AlphaSLiM score, defined as:

*AlphaSLiM score = N × (pLDDT_bound - pLDDT_unbound)*

where N is the PLIP-derived interaction count. Motif regions were defined as contiguous residues with positive pLDDT differences, scored by their maximum AlphaSLiM value. While MiniPAE offers simpler implementation, AlphaSLiM remains available:

https://github.com/martinovein/AlphaSLiM.

We also used **MinD**, a metric described by Omidi *et al*. (accessible at https://github.com/alirezaomidi/AFminD), and the **Average_model score**, developed by Schmid *et al*. available at (9).

To facilitate comparative visualization, all scores (AlphaSLiM, MiniPAE, and MinD) were normalized to a 0–1 scale when indicated. AlphaSLiM scores were normalized by division with the maximum observed value. For MiniPAE and MinD, normalization was performed by first taking the inverse of each value, subtracting the minimum inverse value, and then dividing the result by the maximum inverse value.

### Quantification and statistical analysis

All computed statistics and ROC curves (Figure 1-2) were done as implemented in the Prism 8.0.02 statistical analysis software (GraphPad Software, Inc., San Diego, CA).

### Plasmids, retroviral and lentiviral constructs

Expression plasmids are presented in **Table S3**. Note that FLAG tagged proteins expressed by these vectors contain 3xFLAG tag which are referred to as FLAG-in the text and figures.

pBLP10 and pMD108 plasmids expressing HA-RSK(WT)-BioID2 (17) and HA-RSK(mutant)-BioID2 were kindly provided by Belén Lizcano Perret and Melissa Drappier respectively.

All “FLAG-GFP-SLiM” expression plasmids were created by inserting hybridized oligo between *Bsu*36I and *Xba*I restriction sites within pMV51 plasmid, a derivative of pTM952 (18,19). SLiMs coding sequences (**Table S3**) were generates by Novopro codon optimization tool (https://www.novoprolabs.com/tools/codon-optimization)

### Cell culture and transfection

HEK293T (20) and HeLa-BLP32 (expressing analog-sentive human RSK1 (16)) were maintained in Dulbecco’s Modified Eagle Medium (Lonza) supplemented with 10% fetal bovine serum (Sigma), 100 U/mL penicillin and 100μg/mL streptomycin (Lonza). All cells were cultured at 37°C in a humidified atmosphere containing 5% CO2.

Cells seeded the day before were transfected using Lipofectamine 2000 (Thermofisher) according to the manufacturer’s instructions. Twenty-four hours post-transfection, the cells were incubated with 5 µM biotin (diluted in culture medium) for an additional 24 hours at 37°C.

For analog sensitive kinase experiments, cells were incubated with analog sensitive kinase buffer prior to lysis, following the protocol described by Lizcano-Perret *et al*. (17).

### Immunoprecipitations and Western blot analysis

Immunoprecipitation and Western blotting were done as previously described (14). Briefly, transfected cells were lysed in lysis buffer (Tris-HCl 100 mM pH 8, NaCl 150 mM, NP40 0.5%, EDTA 2 mM, and supplemented with protease/phosphatase inhibitors (Pierce)) and centrifugated at 14,000 x g for 10 min at 4°C. Cleared supernatants were incubated with anti-FLAG M2 Magnetic Beads (#M8823, Sigma-Aldrich) with gentle agitation for 4 hours at 4°C. Magnetic beads were then washed 3 times with the lysis buffer. Immunoprecipitated proteins were detected by Western-blot analysis antibodies listed in Table S3.

## Data availability

All data are available for download and biological material is freely available upon request. All benchmark Alphafold2 data (ELM_dataset including noPDB_dataset, ∼200Gb in 5 parts) is available at: https://zenodo.org/uploads/15260141, https://zenodo.org/uploads/15260224,

https://zenodo.org/uploads/15273252, https://zenodo.org/uploads/15273254, https://zenodo.org/uploads/15273263.

The monomeric predictions additionally required for AlphaSLiM are available at: https://zenodo.org/uploads/15236595.

AlphaFold3 predictions for noPDB_dataset are available at: https://zenodo.org/uploads/15236655.

hRSK2 SLiM predictions are available at https://zenodo.org/uploads/15235709. The monomeric predictions required for AlphaSLiM are available at: https://doi.org/10.5281/zenodo.15236154

## Discussion

Although SLiMs regulate nearly all biological processes, they remain notoriously difficult to detect, both *in-vitro*—due to their often low-affinity interactions—and *in-silico*, because of their short size and degenerate sequence nature.

To address this challenge, recent advances in AI-based protein structure prediction, particularly AlphaFold, have opened promising avenues. In this study, we focused on using different AlphaFold2 and AlphaFold3 scoring functions to detect SLiMs independently of the overall structural quality, allowing us to explore motifs that are entirely absent from PDB, which contrasts with prior approaches reliant on structural templates.

To do so, we generated a benchmark dataset derived from the ELM database and showed that MiniPAE, computed on AlphaFold2 predictions, performs comparably to both MinD and our AlphaSLiM scoring function in detecting SLiMs. While all three scores are effective, its simpler implementation makes MiniPAE the most practical option for general users: it does not require Alphafold extra output that drastically increases prediction size (as minD does), nor does it require additional monomer predictions (as AlphaSLiM does). These conclusions are specific to SLiM prediction and may not extend to other types of IDR-mediated interactions, as previously suggested by Omidi *et al.* (6).

Notably, MiniPAE scores derived from AlphaFold3 predictions were slightly lower than those from AlphaFold2, aligning with recent findings comparing the two models for peptide–protein interactions (21). Given this performance difference—along with the current limitations in prediction throughput on the AlphaFold server—AlphaFold2 remains the more practical choice for this application.

On the experimental side, high-throughput phage display screening of SLiMs was performed for the first time in December 2021, achieving a recall of 23%. However, the prohibitive cost and the tremendous workload of protein purifications of this technique limits its adoption. In contrast, our *in-silico* workflow offers a recall of 30% and at a relatively low cost. That said, AlphaFold’s limited specificity currently precludes blind, proteome-wide screening (5,9). While this may appear restrictive at first, once a SLiM consensus has been identified, it can be used to search for new potential partners using dedicated sequence-based tools such as the widely adopted SLiMSearch4 (10). Alternatively, to reduce specificity limitation and make a broader screen, users can re-score their predictions with SPOC (9), which makes use of AlphaFold-independent data. However, SPOC is currently limited to human reference proteins.

Finally, to illustrate the power of our workflow, we applied it to the RSK2 kinase. Although many of the predicted motifs were homologous to known SLiMs, we identified 14 previously undescribed RSK2-binding SLiMs. Using our highly sensitive proximity labelling system, we confirmed to various extents all the tested predictions experimentally. Among these, we found that GSTE1 interacts with RSK2 via two distinct SLiMs: a substrate-like motif divergent from the canonical phosphorylation site, and a DDVF motif, which likely engages a different region near the kinase active site. We hypothesized that this dual SLiM strategy may facilitate the phosphorylation of non-canonical substrates, expanding the functional repertoire of the kinase.

The rise of ColabFold, a Google Colab-powered implementation of AlphaFold2, has democratized the use of structure prediction, allowing any researcher to perform interaction predictions without the need for local installation. Building on this, we provide a Google Colab notebook for SLiM identification across multiple AlphaFold predictions—eliminating installation barriers entirely. Together, these tools aim to make AlphaFold-based SLiM prediction accessible even to researchers without coding expertise.

## Acknowledgements

We thank Alireza Omidi for the original miniPAE script (6), Belen Lizcano-Perret for BLP32 cells, the BLP10 plasmid, and Figure 5A’s kinase illustration (16,17), Melissa Drappier for the MD108 plasmid, Laurent Gatto for benchmarking guidance, and Nicolas Papadopoulos for advice on AlphaFold screenings. We used ChatGPT to support code writing and language editing.

## Supplementary material

**Table S1 - Benchmark for AlphaFold SLiM predictions**

**Table S2 - RSK2 SLiM prediction**

**Table S3 - Plasmids & Antibodies**

**Figure S1:**
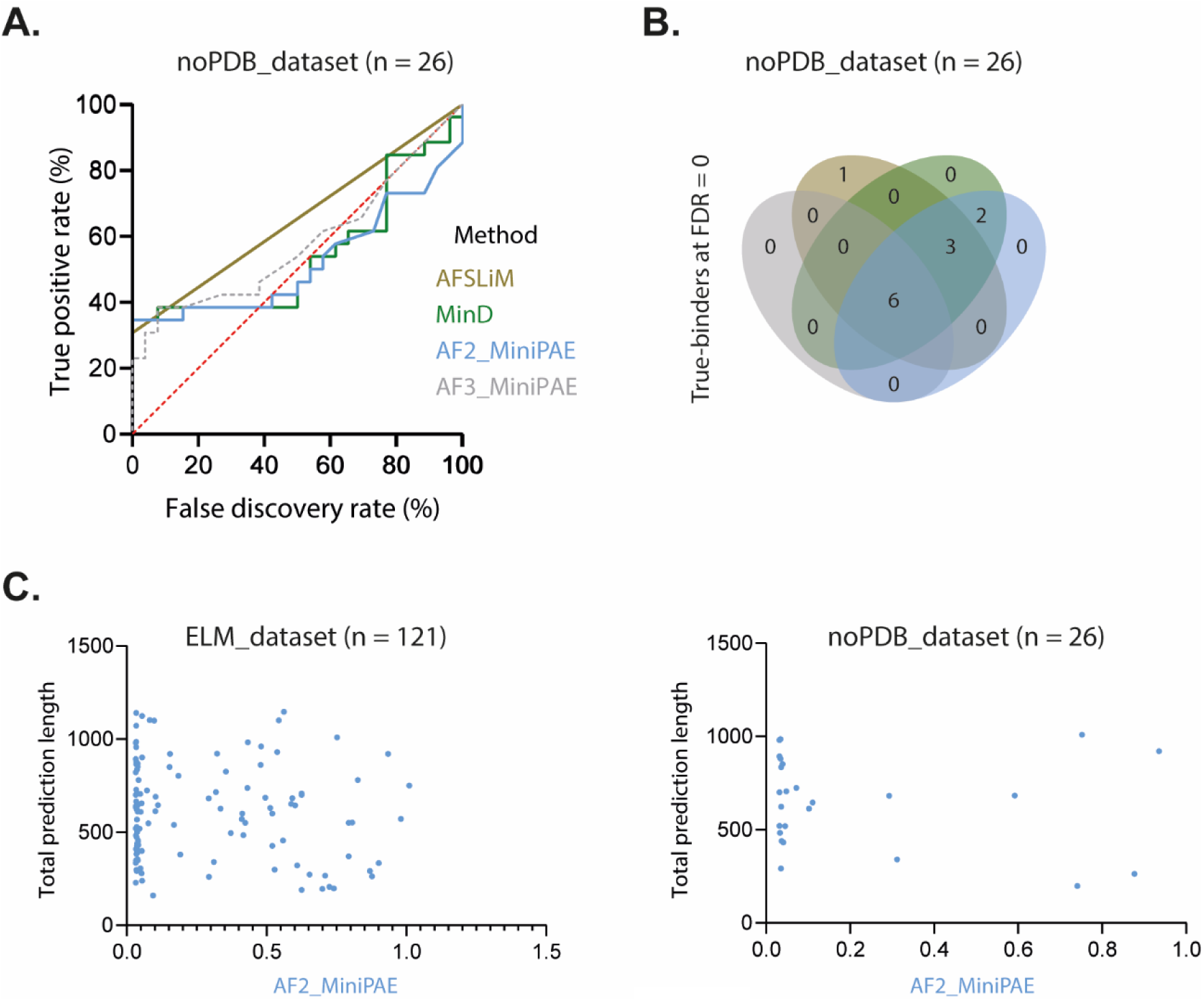
Additional ways to analyze SLiM discovery by AlphaFold within fulllength proteins. **A.** ROC curves characterizing AlphaFold-based scores in the SLiM region when paired with true-binders or a non-binder. **B.** Venn diagram comparing different scoring functions. **C.** Absence of correlation between total prediction length and AF2_MiniPAE score. AF2/3 = AlphaFold2/3. AFSLiM = AlphaSLiM.

